# Why high yield QTLs did not succeed in preventing yield stagnation in rice?

**DOI:** 10.1101/634196

**Authors:** Dongliang Xiong

**Author notes:** **Corresponding author**: Dr. Dongliang Xiong, or.

## Abstract

Rice plays a vital role in global food security, and its yield needs to be increased to meet escalating demand. Although many high yield quantitative trait loci (QTLs) have been identified in the last decades, rice grain yield in the main rice-producing countries is stagnating since the middle of the 1990s. By summarizing the yield performance of high-yielding QTL lines, we found that almost all the high-yielding QTL introduced lines had no practical usage in current high yield breeding programs, due mainly to their low absolute grain yield. Further analysis showed that scientists primarily focused on spikelets number per panicle alone rather than other yield traits, and, in most of the studies, the yield increase was referenced to very old cultivars. By analyzing the yield traits correlations across cultivars in both field and pot conditions, and yield traits correlations across different eco-sites using the same cultivars, we demonstrated that the rice high yield will be rarely achieved by using single-trait approaches due to the traits trade-offs. Building on this, several recommendations are provided to the next generation of biotechnological breeding in rice.

## 1. Introduction

Enhancing crop production is a global challenge since the rate of population growth currently exceeds the increase of food production (Godfray et al., 2010). It has been forecasted that more than 70% primary foodstuffs will be needed by 2050 (Long et al., 2015). Because of rapid urbanization process, available land for crops cultivation has been decreasing rapidly; therefore, increasing grain yield per unit land area is envisaged as the only way to meet the increasing global demand (Long et al., 2015). Agricultural producers draw on a multitude of technologies to ensure efficient, sustainable, stable and high-quality crop production. Genetics has been the foundation of crop improvement since the dawn of agriculture. New genome technologies have transformed genetics into an information-rich (driven) discipline.

Rice (*Oryza sativa* L.) is one of the most important crops for more than half of the world’s population, and its grain yield is determined by four yield components: number of panicles per unit area, spikelet number per panicle, seed setting rate, and grain weight. Theoretically, improving rice grain yield can be potentially achieved by increasing a single yield component or any combination of multiple increases. However, the seed setting percentage in rice is very close to its upper limit, and its further improvement might be limited. Rice genome has been well sequenced, and structural and functional rice genomics have been analyzed based on its full genome sequence (Goff et al., 2002; Yu et al., 2002; Wang et al., 2018). Such studies facilitated marker development and quantitative trait loci (QTLs) identification (Xu et al., 2012; Chen et al., 2014). Over the last decades, many high-yielding rice QTLs have been identified (Xing and Zhang, 2010; Miura et al., 2011; Wing et al., 2018), such as *GN1a* (Ashikari et al., 2005), *GhD7* (Xue et al., 2008a), *SPIKE* (Fujita et al., 2013) and *TGW6* (Ishimaru et al., 2013), and it has been reported that grain yield increased over 50% under experimental conditions by integrating a single QTL of *Ghd8* (Yan et al., 2011). Moreover, high yield varieties are expected to be achieved by pyramiding high-yielding QTLs using biotechnology more rapidly compared to conventional breeding (Long et al., 2006; Xing and Zhang, 2010). However, the average annual rice grain yield in China and Japan - where most of high yield QTLs were identified – are still stagnating (Fig. S1). Facing this picture, one may rightfully ask: why identifying high yield QTLs (genes) did not dramatically improve rice grain yield in farmland? If the biotechnological strategies did not work in the near future, how should the further 70% additional rice yield be obtained?

Here we performed a synthesis analysis based on published data, new data generated by this study, and a comparative study between Taoyuan and other yield testing nurseries to reveal why high yield QTLs did not work well in improving rice yield.

## 2. Materials and Methods

### 2.1 High yield QTLs

To evaluate the contributions of high yield QTLs to rice grain yield under non-stress conditions, research papers were searched from the Web of Science database (Clarivate Analytics, Boston, MA; Table S1). To date, hundreds of QTLs for rice yield have been reported, however, only a few of these QTLs have been clearly characterized. To ensure the conclusions reliable, only the clearly characterized QTLs (genes) were included. The studies without grain yield of the wild-type and/or its QTL introduced mutants were excluded from our analysis. Hence, some reputed high yield QTLs, such as *GN1A* (Ashikari et al., 2005), *GhD7* (Xue et al., 2008b), *GS5* (Li et al., 2011) and *DTH8* (Wei et al., 2010), had to be excluded due to the lack the grain yield values. Moreover, the high yield QTLs under abnormal growth conditions (i.e. stress conditions) were also excluded. Grain yield of both wild-type and its QTL introduced mutants were extracted as well as yield components when available.

### 2.2 Field experiment

Four field experiments were conducted at Wuxue, Hubei, China (29°59’50”N, 15°36’56”E) in 2013-2015. The soil chemical properties were shown in Table S2. Totally, 55 genotypes with high variable spikelets number per panicle as well as panicle number were used in those experiments. A completely randomized block design with four replications was adopted in all the experiments, and rice plants were grown in different growth seasons to create the variability. Local field management was adopted. At physiology maturity, 12 hills in each plot were harvested to determine the yield components. Panicle number was converted to per square meter multiplying by plants’ density. The filled spikelets were separated from unfilled spikelets after submerging them in tap water, and the empty spikelets were separated from the half-filled spikelets through winnowing. The spikelets per panicle, and grain filling percentage (100 × filled spikelet number/total spikelet number) were calculated. The grain yield was determined from a 5 m^2^ area in each plot (30 m^2^). For all the measurements of four field experiments were following the same protocol.

### 2.3 Pot experiment

The pot experiment was conducted in Huazhong Agriculture University, Wuhan, China in 2014. In this experiment 88 genotypes were used. Seeds were sow in 13.0 l pot filled with paddy soil and four replicates for each genotype. At two leaves stage, the plants were thinned to single plant per hill with a density of three hills per pot. During the experiments, the plants were well watered (at least 2cm water level was maintained). Pests and diseases were controlled using insecticides and fungicides. At physiological maturity, yield and yield components for each pot were measured. The measurements of spikelets number per panicle, seed setting percentage, and grain weight were performed following the same protocol as in the field experiments. The grain yield and panicle number were measured for each separate pot.

### 2.4 High rice yield in Taoyuan

The peer reviewed articles were searched from the Web of Science, Scopus, and the China Knowledge Resource Integrated Database using three search terms: ‘Rice AND Taoyuan’, ‘Rice AND Yongsheng’, and ‘Rice AND Yunnan’. The data of grain yield and yield components were extracted directly from the tables in the original papers, or indirectly from figures using WinDIG 2.5 (http://www.unige.ch/sciences/chifi/cpb/windig.html). Other information, if available, such as experimental sites location, and cultivar name were also extracted for further analysis. The summary information of those studies can be found in Table S3.

### 2.5 Statistical analysis

One-way ANOVA analysis was used to test the differences in yield and yield component traits at Taoyuan and at other sites. Regression analyses were performed to test the correlations between parameters. All analyses were performed in R (R Core Team, 2018).High yield QTLs

## 3. Results

### 3.1 High yield QTLs

In our database, 19 identified rice high yield QTLs were included (Table S1, Fig. 1A) and most of the QTLs are related to large panicles. Overall, high-yielding QTLs contribute differently to the realized grain yield under experimental conditions (Fig. 1A, Fig. S2), with a range of from 1.14% (*gw7*) to 70.8% (*Ghd* 8). Comparing with their respective wild types, the plants with a single high-yielding QTL showed increased grain yield by 22.01 *%* on average. Grain yield of plants with QTLs for spikelets number per panicle, seed setting percentage, grain weight and panicle number increased by 29.51%, 6.17%, 3.78% and −0.58%, respectively (Fig. 1B, Fig. 2).

**Fig. 1.**
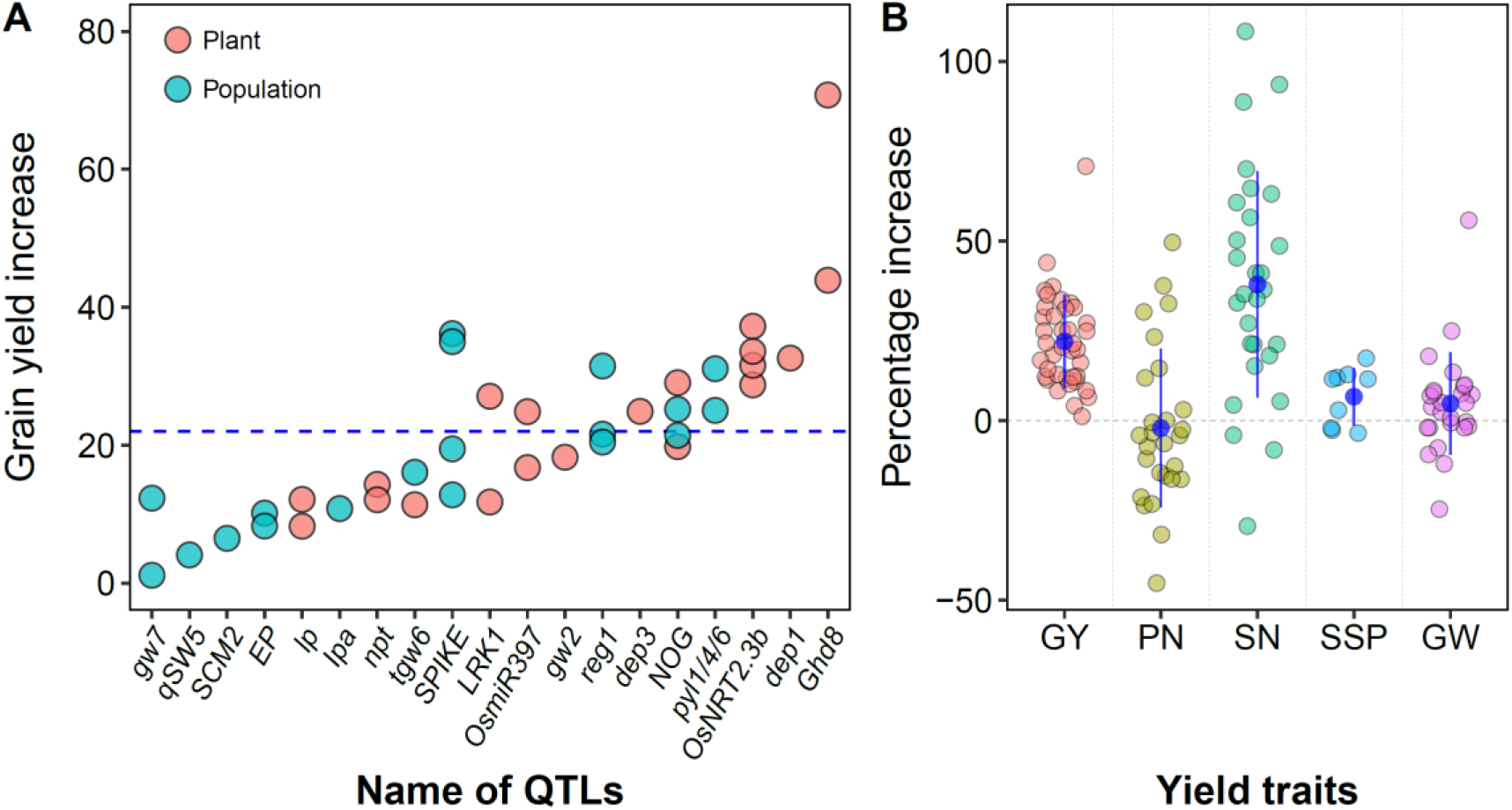
Contributions of high yield genes on yield increase (A) and average increase of yield and yield components in high-yielding QTL introduced lines (B). The dot line in (A) represents the average grain yield increase (22.01 %), and the blue points and bars in (B) represent the means and standard errors. GY, Grain yield; PN, Panicle number; SN, spikelets number; SSP, seed setting percentage; GW, grain weight. The percentage increase was calculated as 100%×((*X*_QTL_-*X*_wild_)/ *X*_QTL_), where the *X*_QTL_ and *X*_wild_ represent the parameter values for plants with introduced QTL and wild type, respectively.

**Figure 2.**
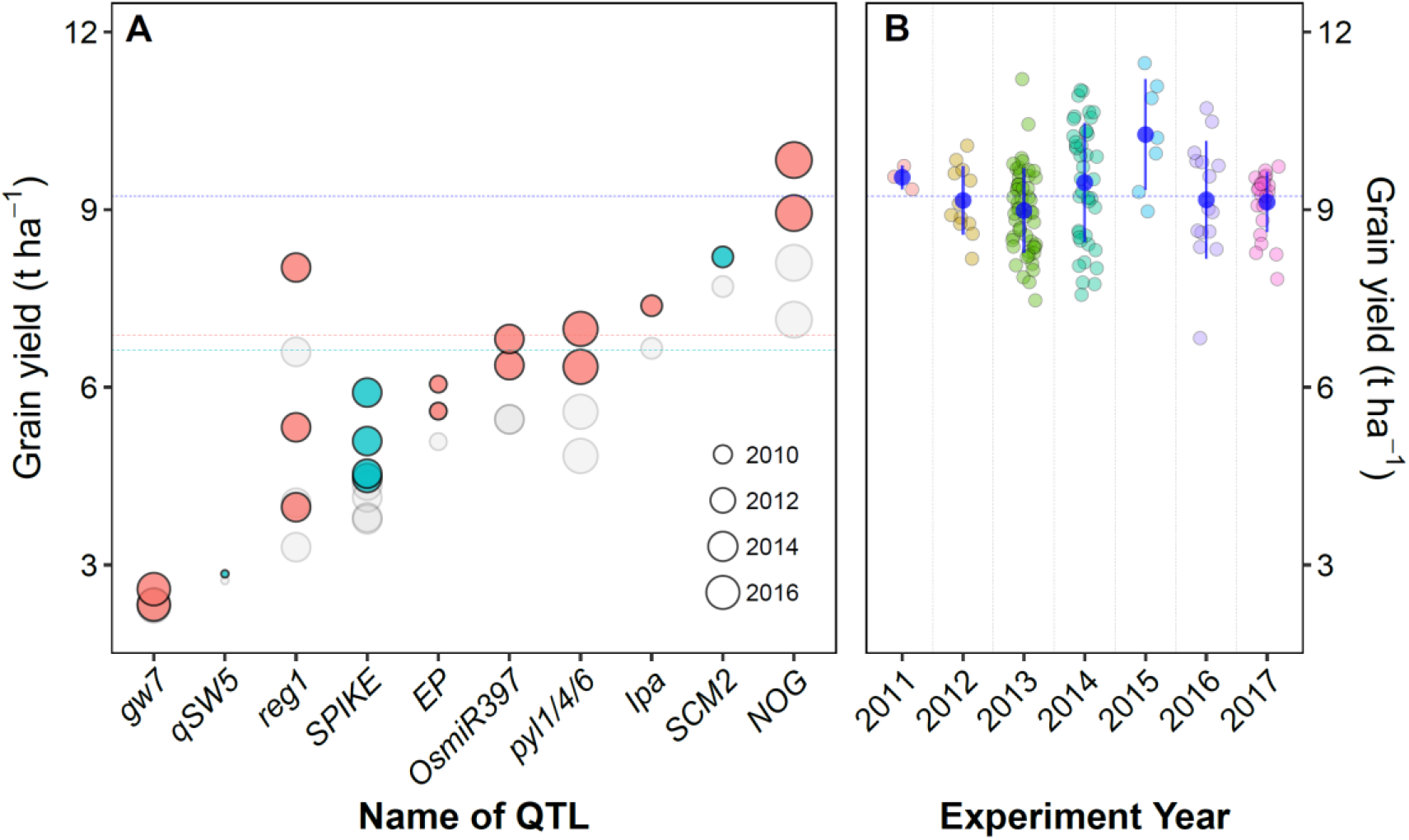
(A) Effects of high yield QTLs on rice grain yield on population level, and (B) grain yield of Yangliangyou 6 (YLY6) - a widely used reference cultivar in promotion trials of newly developed varieties in China - under field condition at Wuxue, Hubei, China. In panel (A), the red and green points represent the high yield QTL identified in China and Japan, respectively, and the grey points represent the wild types (references). The red and green dot line represent the national average annual grain yield of China and Japan in 2015 (data from FAO), respectively. The blue dot line in both (A) and (B) represents the average grain yield of YLY6 under field condition in over seven years at Wuxue, Hubei, China.

### 3.2 Panicle number vs spikelet number

Under both field (Fig. S3) and pot (Fig. S4) conditions, panicle number, spikelet number per panicle, seed setting percentage and grain weight varied significantly across the dataset. As the result, rice grain yield ranged from 4.0 t ha^−1^ to 9.84 t ha^−1^ under field condition, and ranged from 20 g pot^−1^ to 140 g pot^−1^ under pot conditions. Under field condition, the correlations between grain yield and single yield component were not clear, however, under pot condition, the grain yield significantly correlated with seed setting percentage (Fig S3-S4). A strong negative correlation between panicle number and spikelet number per panicle was observed under both field and pot conditions (Fig. 3).

**Fig. 3.**
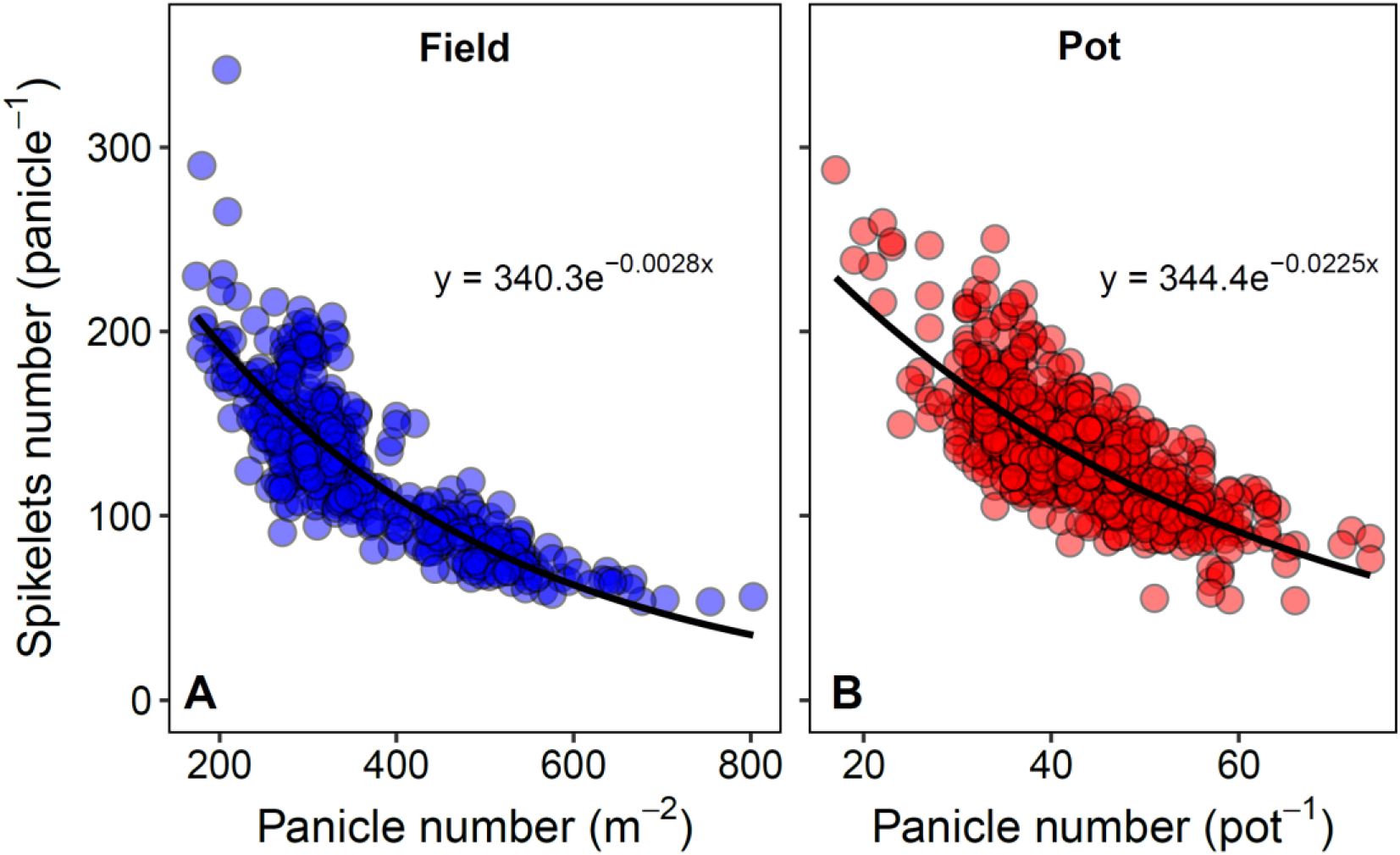
The correlation between spikelet number per panicle and panicle number across rice genotypes under field (A) and pot (B) conditions.

### 3.3 Rice yield in the special eco-site Taoyuan

The average grain yield at Taoyuan was significantly higher than that of the sites outside Taoyuan (Fig. 4). The average grain yield in Taoyuan was 14.2 t ha^−1^ with the ranges from 6.3 t ha^−1^ to 18.7 t ha^−1^ and the average grain yield of other sites was 9.07 t ha^−1^ with the ranges from 5.5 t ha^−1^ to 13.2 t ha^−1^. As expected, the yield components, except grain weight at Taoyuan were significantly higher than the average of other sites. However, the correlations between four yield components and grain yield at Taoyuan or across other sites were not clear (Fig. S5). The negative correlation between panicle number per area and spikelet number per panicle was observed at both Taoyuan and the across other sites (Fig. 5). However, the under a given panicle number per area, the spikelet number per panicle at Taoyuan is higher than other eco-sites.

**Fig. 4.**
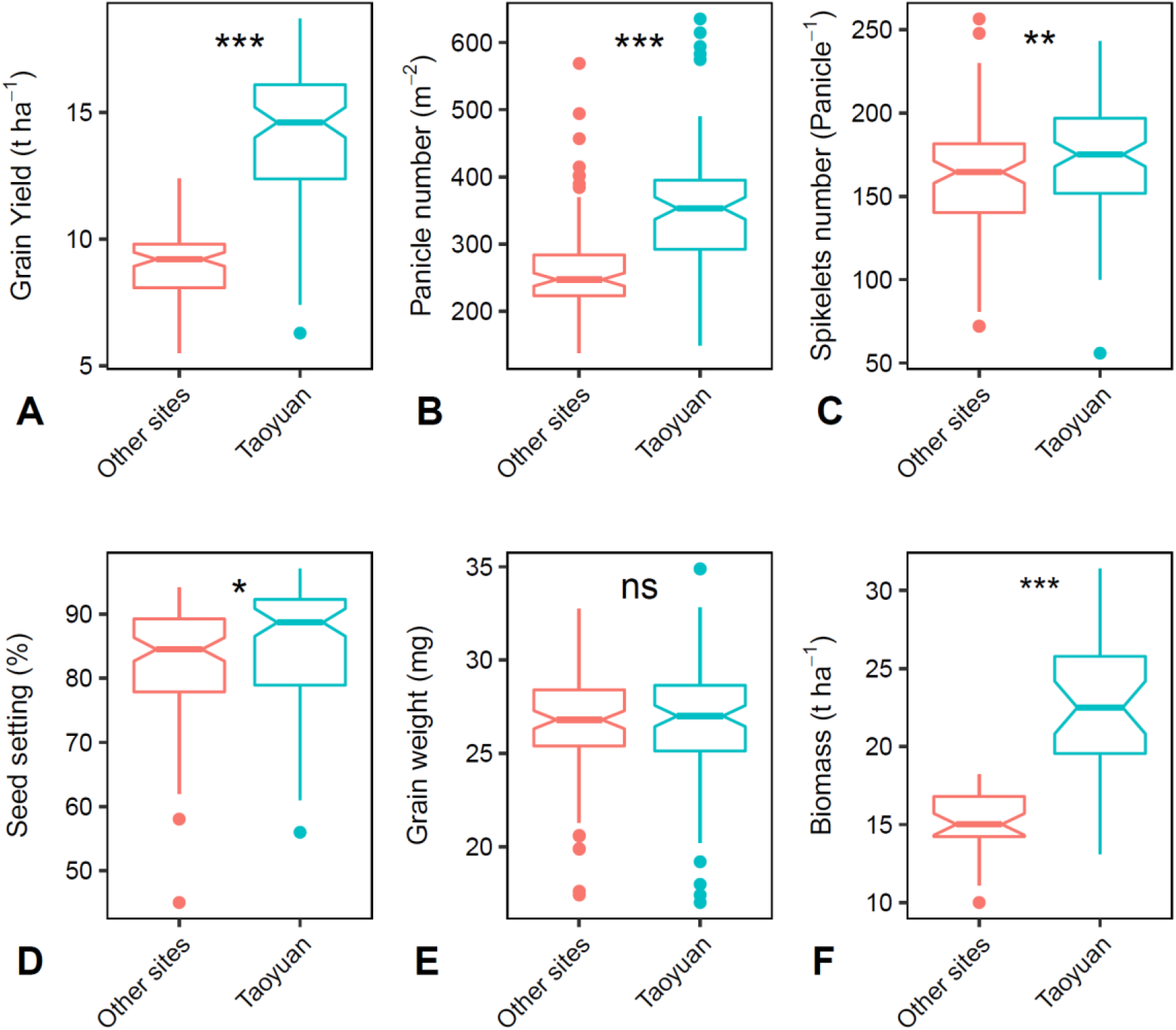
Rice yield and yield components in a high yield eco-site, Taoyuan, and other eco-sites. *, p<0.05; **, p<0.01; ***, p<0.001; ns, p>0.05.

**Fig. 5.**
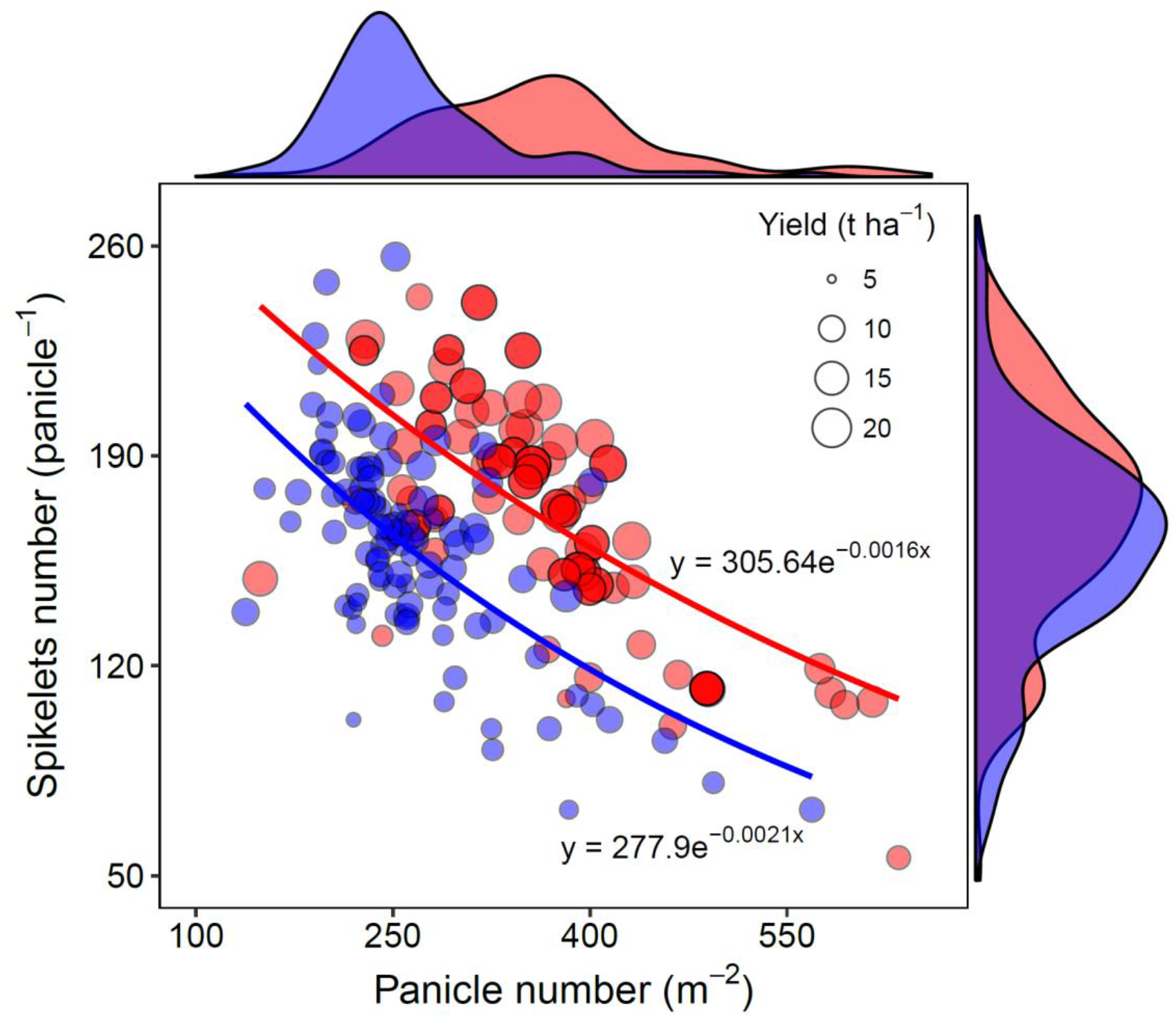
Correlation between spikelet number per panicle and panicle number per unit land area. The blue and red points represent rice grown at other sites and at Taoyuan, respectively.

## 4. Discussion

We constructed a dataset including agronomic traits of high-yield QTL lines and their wild-types reported by rice geneticists over past few decades (Fig. 1; Supplementary table 1). In some of these studies, the grain yield was expressed on a plant level basis (i.e. g plant^−1^) rather than on population level (i.e. t ha^−1^) which was practiced widely, and to make grain yield comparable, the lines that grain yields were expressed on population level were selected for further comparing analysis (Fig. 2). Surprisingly, the rice grain yields of many high-yielding QTL introduced lines were very low (range from 2.33 to 9.84 t ha^−1^, with an average of 5.88 t ha^−1^), and in most of these lines, the grain yield was even lower than the national annual grain yield (2015) in China and Japan, in where these QTLs were identified. In fact, the enhancements of high-yielding QTL lines were based on ancient rice cultivars that have low yield potential, for instance, Nipponbare (released in 1963) and Zhonghua 11 (released in 1979) in these studies. Disappointingly, only one high-yielding QTL introduced line (*NOG* with background of Teqing) has the grain yield close to the average grain yield of YLY6, a widely used reference cultivar in promotion trials of newly developed varieties in China (Fig. 2B), which indicates that these high-yielding QTL introduced lines might have no practical usage in high yield breeding, as grain yields of newly developed rice cultivars are often higher than 10.0 t ha^−1^ in experimental conditions since 2000 in both China and Japan (Njinju et al., 2018). One possibility is that some of these high-yielding QTLs have been introduced to the modern rice cultivars unconsciously by breeders; however, we cannot test this hypothesis in our current analysis, and further investigations are needed.

Another interesting picture in our analysis is that most of the high-yielding QTLs related to spikelet number per panicle rather than other yield components (Fig. 1). As shown in Fig 1B and Fig. S2, most of the high-yielding QTL introduced lines increased spikelet number per panicle, except the *gw7* and *gw2* in which the enhancement of grain yield was mainly through increase grain weight. Hence, we may ask: can rice grain yield be enhanced by increasing spikelet number per panicle alone? To test this hypothesis, we firstly analyzed the yield traits correlations across genotypes with different panicle size and panicle number under both field and pot conditions. No significant correlation between grain yield and any single yield component was observed, except a weak correlation between grain yield and spikelets number under field condition and between grain yield and seed setting percentage in pot condition. It should be pointed out that the extremely low seed setting percentage of some genotypes in pot condition (Fig. S4) was intruded by high temperature during the flowering. Our results indicate that the variation of grain yield is not determined by a single yield component trait.

A negative correlation was observed between panicle number and spikelets number per panicle under both field and pot conditions (Fig. 2), which indicated the existence of a strong trade-off between these two traits. Considering the grain yields of more than half of the high-yielding QTL lines were reported based on single plant, the effect of increasing spikelets number per panicle on grain yield may be offset by the reduce of panicle number at the population levels. The mechanism for the counteracting effects can be illustrated by the sink – source balance theory: in a unit space, less panicle number may have the benefit for individual panicles to accumulate more carbon during the spike development, which can support more spikelets per panicle; in contrast, more panicle numbers may increase the competition among panicles, and then reduce the investment to spikelets on each panicle. Assuming that seed setting and grain weight can be maintained, further enhancement of grain yield requires an increase of spikelets number per panicle and panicle number per land area simultaneously, which likely depends on an improvement of rice biomass accumulation capacity per land unit.

To validate the statement, a synthesis analysis was performed to compare the rice yield formation of same genotypes in Taoyuan, a well-known rice high-yield eco-site because of its favorable ecological conditions (Ying et al., 1998; Katsura et al., 2008) and in other experiment sites (Table S3). As expected, an increase of all the yield components, except grain weight, contributed to the high yield at Taoyuan (Fig. 4, Fig. S6). These results, again, suggest that it is necessary to improve multiple yield components synchronically, instead of improving spikelet number per panicle alone, to achieve a higher grain yield. Importantly, the negative correlation between panicle number and spikelets number was observed in Taoyuan and across other sites. This confirms the counteracting effects of panicle number and spikelets number. Although the seed setting percentage in Taoyuan (85.2 %) was slightly higher than in other sites (82.3%), the higher grain yield at Taoyuan can be largely explained by the shift of correlation between panicle number and spikelet number per panicle (Fig. 5). Further breeding and planting managements for high rice yield should, therefore, target at reducing the counteracting effects of panicle number and spikelets number per panicle.

## 5. Conclusion and perspectives

Here we show that almost all the high-yielding QTL introduced lines had no practical usage in current high yield breeding practices, due mainly to their low absolute grain yield in the field. Several candidate reasons may contribute to the limited success in achieving high yield by introducing high-yielding QTLs. Single-trait, specifically, the spikelets number per panicle, approaches have been used in most cases, but rice yield is an extremely complex trait that is determined by multiple components as showed in this study. Moreover, the counteracting effects existing between panicle number per land area and spikelets number per panicle further confirmed that single trait approaches may hard to match the high yield breeding goal in rice. These results suggest future research should focus on multiple traits approaches. Fortunately, the multiple traits approach has begun to be concerned, for instance, a very recent research showed that rice grain quality of a high yield potential cultivar can be significantly improved without yield loss by introducing multiple grain quality genes (Zeng et al., 2017).

Another reason may be that the high-yielding QTL lines in most of the studies were created by introducing high-yielding QTLs to very old rice cultivars that usually have low yield potential. Although studies that investigate whether some of these high-yielding QTLs have been introduced to the modern rice cultivars unconsciously by breeders are still needed, we recommend researchers use modern cultivars as backgrounds or at least add some modern cultivars as really high yield references. Importantly, to take the counteracting effects into account and to move to the actual production practices, evaluating the contributions of high-yielding QTLs to grain yield on the population level rather than plant or panicle level is highly recommended.

## Acknowledgments

This work of JH is supported by a grant from the National Key Research and Development Program of China (No. 2016YFD0300210) and a grant from the National Natural Science Foundation of China (No. 31671620). The work of JF is supported by Ministerio de Economía y Competitividad (Spain, Plan Nacional project CTM2014-53902-C2-1-P). The authors wish to thank Tingting Yu, Dan Wang and Xuewu Zhan for their helps on the field and pot experiments data collections and Dr. Fei Wang for helpful comments.

## Supplementary information

**Table S1**. The high yield QTLs of rice analyzed in the current study.

**Table S2**. The soil chemical properties at Wuxue, Hubei, China.

**Table S3**. Summary information of comparing experiments at Taoyuan and at other sites.

**Fig. S1** The average annual grain yield of rice per unit land area. The lines are the loess best fit to the trend of yield against time. Data from FAO (2016).

**Fig. S2** effects of high yield QTLs on rice grain yield (GY), panicle number (PN), spikelets number (SN), seed setting percentage (SSP) and grain weight (GW). The percentage increase was calculated as 100%×((*X*_QTL_-*X*_wild_)/ *X*_QTL_), where the *X*_QTL_ and *X*_wild_ represent the parameter values of plants introduced QTL and wild type, respectively.

**Fig. S3** the correlations among grain yield (GY), panicle number (PN), spikelets number (SN), seed setting percentage (SSP) and grain weight (GW) under field condition.

**Fig. S4** the correlations among grain yield (GY), panicle number (PN), spikelets number (SN), seed setting percentage (SSP) and grain weight (GW) under pot condition.

**Fig. S5** The correlations between grain yield and yield components. The blue and red points represent at Taoyuan and other sites, respectively.

