## Supplementary material for "Why high yield QTLs did not succeed in preventing yield stagnation in rice?": supp materials

Dongliang Xiong*, Shaobing Peng, Jianliang Huang, Cyril Douthe, and Jaume Flexas

**Corresponding author:**

Dr. Dongliang Xiong

College of Plant Science and Technology, Huazhong Agricultural University, Wuhan, Hubei 430070, China

### Supplementary

**Table S1**. The high yield QTLs of rice analyzed in the current study.

| QTL name | Traits | Allele for high yield | Growing condition | Yield expressed | Wild type (release year) | Reference |
|---|---|---|---|---|---|---|
| *DEP1* | Grain number | Loss function | Field | Plant | Zhefu802 (1983) | Huang et al. (2009) |
| *DEP3* | Panicle size; grain shape | Loss function | Field | Plant | Hwacheong (1972) | Qiao et al. (2011) |
| *EP* | Erectpose panicle | Over expression | Field | Population | Liaojing (2001) | Wang et al. (2009) |
| *Ghd8* | Delayed flowering | Over expression | Field | Plant | Zhenshan97 (1968)<br>9311 (1997)<br>Nipponbare (1963) | Yan et al. (2011) |
| *GW2* | Grain size and filling | Loss function | Field | Plant | Fenaizhan1 (1997) | Song et al. (2007) |
| *GW7* | Grain size | Loss function | Field | Population | Huangjingxian74 (2002)<br>Taifeng (NA) | Wang et al. (2015) |
| *IPA* | Grain number; panicle number | High expression | Field | Population/Plant | Xiushui11 (1983) | Jiao et al. (2010) |
| *LP* | Panicle size | Loss function | Field | Plant | Zhonghua11 (1979) | Li et al. (2011) |
| *LRK1* | Panicles number and size; grain weight | High expression | Green house | Plant | 9311(1997) | Zha et al. (2009) |
| *npt* | Panicle number, size | Loss function | Field | Population | Zhonghua11 (1979) | Wang et al. (2017) |
| *NOG* | enoyl-CoA hydratase/isomerase | High expression | Field | Population | Zhonghua17(2004)<br>Teqing (1984) | Huo et al. (2017) |
| *OsmiR397* | Grain size | Over expression | Field | Population | Zhonghua11 (1979) | Zhang et al. (2013) |

| | | | | | | |
|---|---|---|---|---|---|---|
| *OsNRT2.3* | cytosolic pH | Over expression | Field | Plant | Nipponbare (1963) | Fan et al. (2016) |
| *pyl1/6* | ABA receptor | Loss function | Field | Population | Nipponbare (1963) | Miao et al. (2018) |
| *qSW5/GW5* | Grain size | Loss function | Field | Population | Nipponbare (1963) | Shomura et al. (2008)<br>Weng et al. (2008) |
| *Reg1* | Panicle size | Loss function | Field | Population | Zhefu802(1983)<br>9311(1997)<br>Nipponbare (1963) | Li et al. (2013) |
| *SCM2/APO1* | Panicle number and size | High expression | Field | Population | Koshihikari(1956) | Ookawa et al. (2010) |
| *SPIKE/ NAL1* | Panicle size | Over expression | Field | Population | IR64(1985) | Fujita et al. (2013) |
| *TGW6* | Grain weight | Loss function | Field | Plant | Nipponbare (1963) | Ishimaru (2003)<br>Ishimaru et al. (2013) |

Note: the references list can be found at the end of file.

Table S2. The soil chemical properties at Wuxue, Hubei, China.

| Year | pH | Organic matter g $kg^{-1}$ | Total N g $kg^{-1}$ | Available P mg $kg^{-1}$ | Available K mg $kg^{-1}$ |
|---|---|---|---|---|---|
| 2013 | 5.47 | 29.10 | 2.2 | 12.14 | 92.2 |
| 2014 | 5.60 | 27.18 | 1.83 | 4.91 | 105.8 |
| 2015 | 5.20 | 16.69 | 1.19 | 22.5 | 159.2 |

Table S3. Summary information of comparing experiments at Taoyuan and other sites.

| Experiment Year | Site | References |
|---|---|---|
| 1995-1996 | Los Banos, Philippines | Ying et al. (1998a); Ying et al. (1998b) |
| 2002-2003 | Kyoto, Japan | Katsura et al. (2008) |
| 2006-2007 | Nanjing, China | Li et al. (2009) |
| 2007-2008 | Nanjing, China | Li et al. (2014a) |
| 2008-2009 | Nanjing, China | Li et al. (2014b) |
| 2010-2011 | Hongjing, China | Xia et al. (2016) |
| 2011-2012 | Danyang, China | Zhang et al. (2014) |
| 1998 | Binchuang, China<br>Hangzhou, China | Yang et al. (2004) |
| 1998 | Longhai, China | Yang et al. (2000a) |
| 1998 | Longhai, China | Yang et al. (2000b) |
| 1989/1993/1995 /1997/1998 | Shaxian/Danyang/Tianchang/Wuhan/ Guidong/Mianyang/Pingjiang/Linan/ Xinyang/Ganyu/Dengmai | Yuan et al. (2000) |

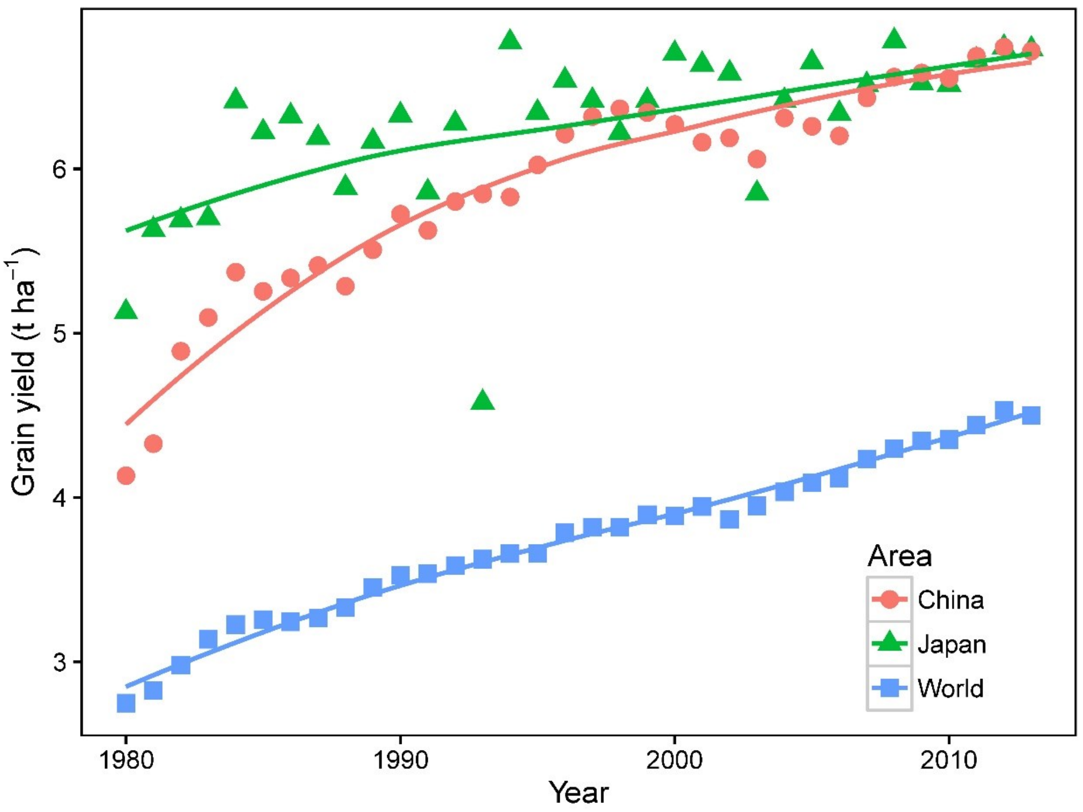


Fig. S1 The average annual grain yield of rice per unit land area. The lines are the loess best fit to the trend of yield against time. Data from FAO (2016).

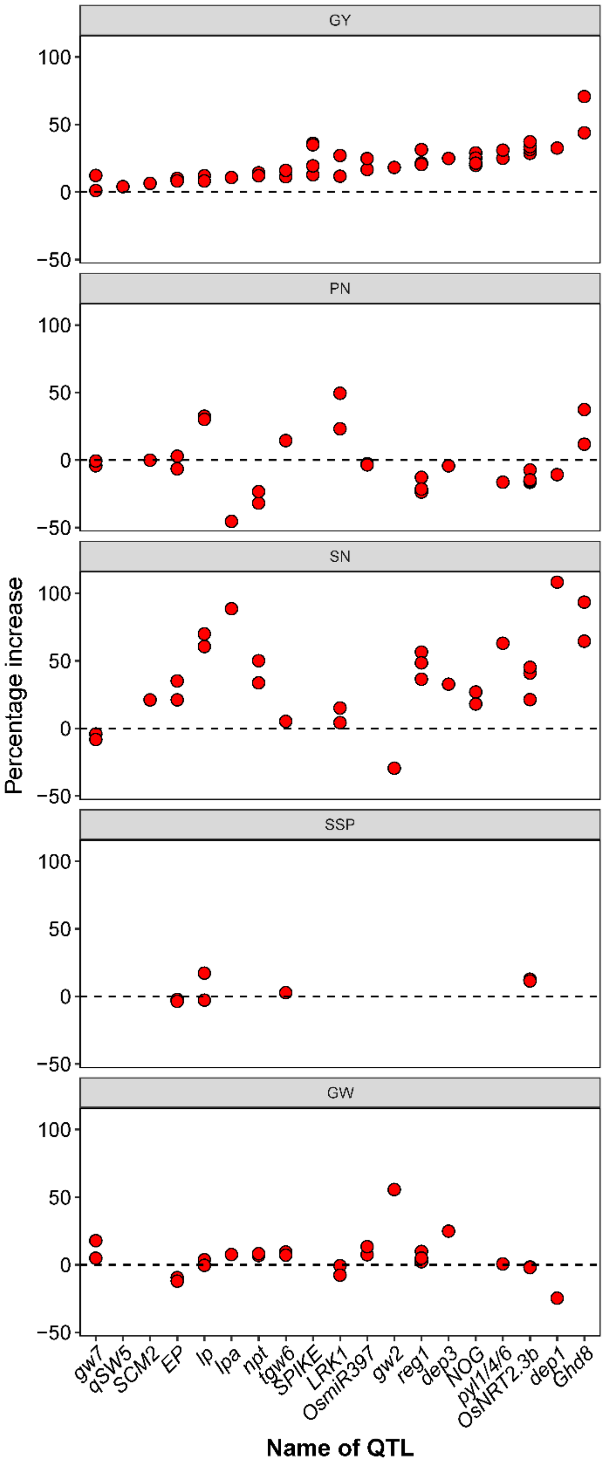


Fig. S2 effects of high yield QTLs on rice grain yield (GY), panicle number (PN), spikelets number (SN), seed setting percentage (SSP) and grain weight (GW). The percentage increase was calculated as 100%×( ( $X_{QTL}$-$X_{wild}$)/ $X_{QTL}$), where the $X_{QTL}$ and $X_{wild}$ represent the parameter values of plants introduced QTL and wild type, respectively.

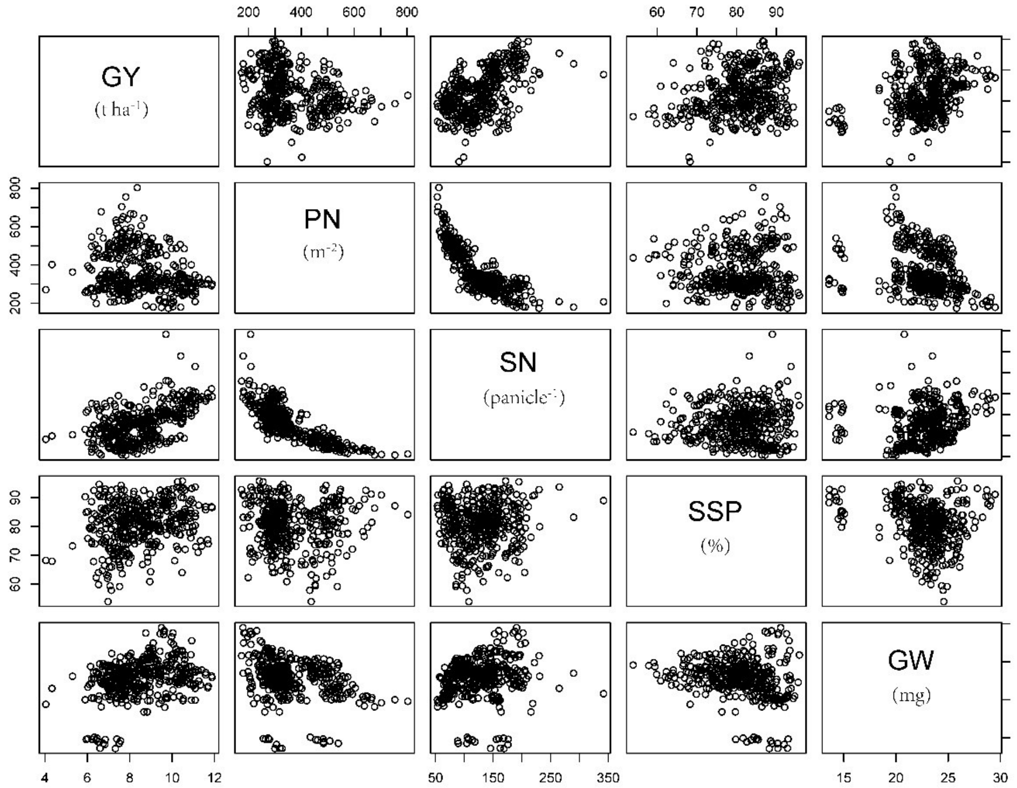


Fig. S3 the correlations among grain yield (GY), panicle number (PN), spikelets number (SN), seed setting percentage (SSP) and grain weight (GW) under field condition.

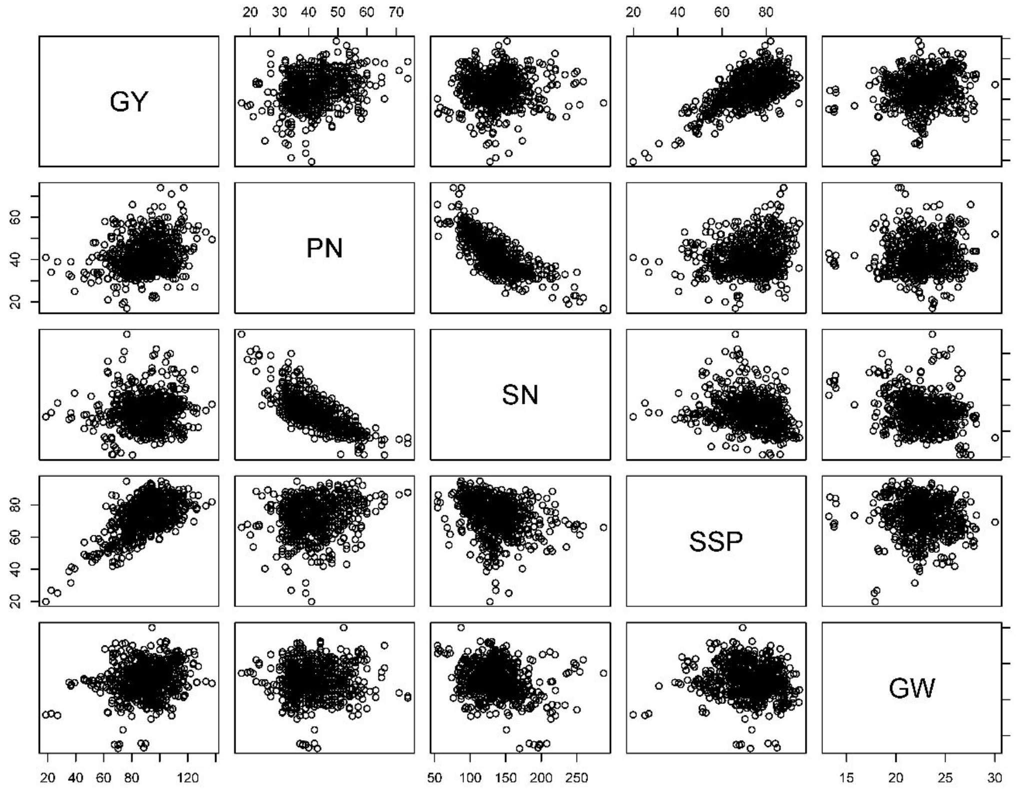


Fig. S4 the correlations among grain yield (GY), panicle number (PN), spikelets number (SN), seed setting percentage (SSP) and grain weight (GW) under pot condition.

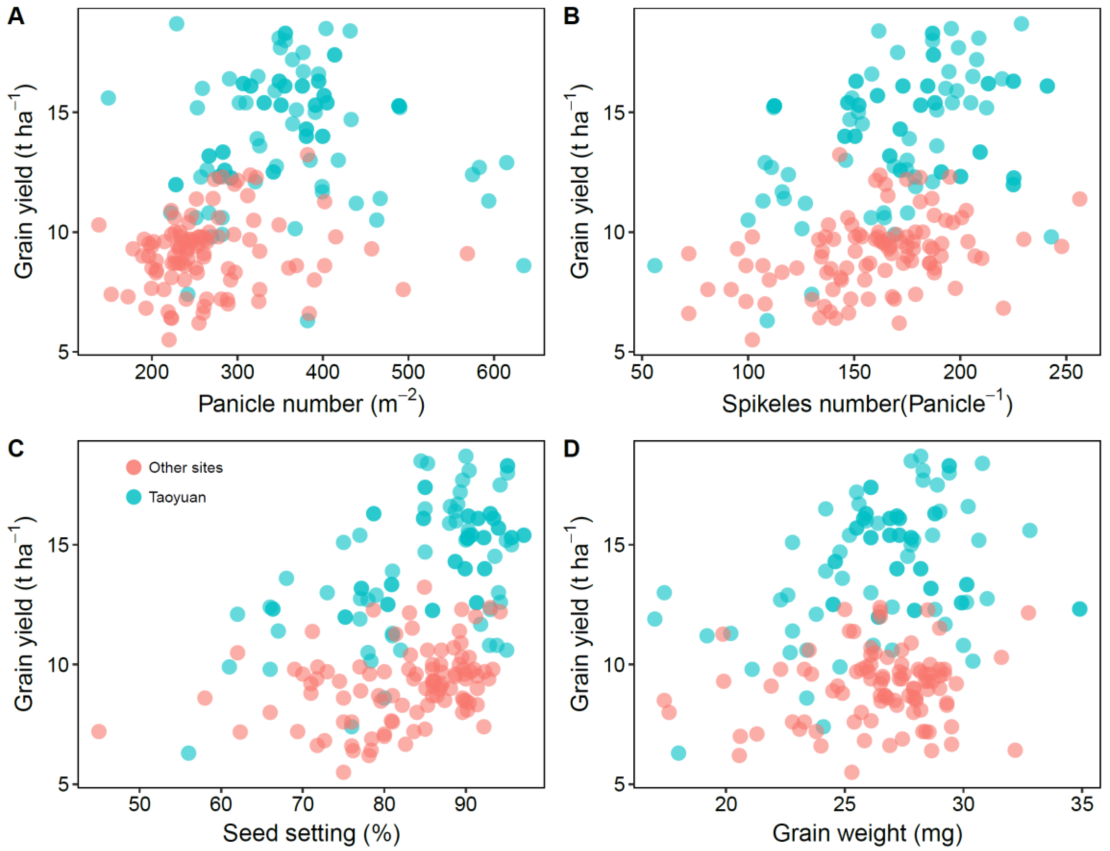


Fig. S5. The correlations between grain yield and yield components. The blue and red points represent at Taoyuan and outside Taoyuan, respectively.
